# Autoregulatory control of microtubule binding in the oncogene, doublecortin-like kinase 1

**DOI:** 10.1101/2020.06.12.149252

**Authors:** Melissa M. Rogers, Amrita Ramkumar, Ashlyn M. Downing, Hannah Bodin, Julia Castro, Dan W. Nowakowski, Kassandra M. Ori-McKenney

## Abstract

The microtubule-associated protein (MAP), doublecortin-like kinase 1 (DCLK1), is highly expressed in a range of cancers and is a prominent therapeutic target for the development of kinase inhibitors. However, the physiological roles of its kinase activity and how DCLK1 kinase activity is regulated remain elusive. Here we employ in vitro reconstitution with purified proteins to analyze the role of DCLK1 kinase activity in regulating microtubule binding. We find that DCLK1 autophosphorylates a single residue within its C-terminal tail to restrict its kinase activity and prevent aberrant hyperphosphorylation within its microtubule-binding domain. Removal of the C-terminal tail or mutation of this residue causes an increase in phosphorylation largely within the doublecortin 2 (DC2) domain, which dramatically reduces the microtubule affinity of DCLK1. Therefore, autophosphorylation at specific sites within DCLK1 have diametric effects on the molecule’s ability to associate with microtubules. Overall, our results suggest a mechanism by which DCLK1 modulates its own kinase activity to tune its microtubule binding affinity, providing molecular insights into a unique form of autoregulatory control over microtubule binding activity within the broader family of MAPs. These results provide useful molecular insights for future therapeutic efforts related to DCLK1’s role in cancer development and progression.

## Introduction

Growth is an essential process of life. Unchecked cellular growth, however, is a hallmark of cancer. Therefore, the process of cell division is often a target of cancer therapeutics (1,2). The macromolecular machine responsible for accurately segregating chromosomes during eukaryotic cell division is the bipolar mitotic spindle, a structure composed of dynamic microtubules organized by a multitude of MAPs (3,4). DCLK1, formerly known as DCAMKL1 and KIAA0369, is one such MAP that is also upregulated in a range of cancers, such as pancreatic, breast, bladder, colorectal, gastric, and hepatocellular carcinoma (5-14). As a consequence, many studies have focused on developing small molecule inhibitors against DCLK1 kinase activity in an effort to control cancer growth (15-17). However, it is currently unclear if DCLK1 kinase activity, microtubule-binding activity, or both are involved in the molecule’s functions during cell division. Therefore, a mechanistic understanding of DCLK1, both at the molecular and biological levels, is currently lacking.

DCLK1 is a member of the doublecortin (DCX) superfamily, which also includes DCX, DCDC2, and retinitis pigmentosa 1 (RP1), all of which are implicated in human disease (15,18-22). At its N-terminus, DCLK1 contains two tandem DCX domains (DC1 or N-DC: aa 54-152 and DC2 or C-DC: aa 180-263)(Figure S1A), which are highly conserved among other family members (6,18,23-25). DCLK1 and its paralogue, DCX, were originally identified and characterized for their functions during neuronal development, including neurogenesis and neuronal migration (5,21,22,26-28). Although the roles of DCLK1 and DCX in neurodevelopment have been phenotypically described in vivo, the molecular basis for these observations remains ill-defined. Prior studies have shown that DCLK1 and DCX may act as microtubule stabilizers, nucleators, and regulators of microtubule-based transport (29-36). Dissecting the mechanisms by which DCLK1 binds to the microtubule can therefore provide insight into the microtubule-binding behaviors of other DCX family members and how they may be subverted in disease.

The C-terminal portion of DCLK1 contains a serine/threonine kinase domain and an unstructured C-terminal tail that share sequence similarities with calcium/calmodulin dependent protein kinase I (CaMKI)(37,38). For both DCLK1 and CaMKI, removal of a distal C-terminal ‘tail’ region results in an increase in kinase activity (37-40). This mode of regulation has been well-studied for CamKI, whose C-terminal tail makes direct contact with the kinase domain, directly inhibiting its enzymatic activity (40). However, it is unclear if, or how, the C-terminal tail of DCLK1 regulates its kinase domain. In addition, the physiological significance of DCLK1 kinase activity is unknown, even though it is a target for the development of kinase inhibitors due to its prominent role in cancer (15-17). Additional information on the functional role of the DCLK1 kinase domain and how it is controlled would therefore be valuable for understanding how drugs can effectively target DCLK1 for therapeutic purposes.

Here we present a detailed examination of the microtubule-binding properties of DCLK1 and how they are regulated by its kinase activity. We find that DCLK1 autophosphorylates one key residue (T688) within its C-terminal tail via an intramolecular mechanism to strongly modulate its microtubule-binding affinity. Removal of the C-terminal tail or mutation of T688 results in an increase in phosphorylation of residues within the DC2 domain, which in turn eliminates microtubule binding. Furthermore, we observe that mutating four key residues within DC2 of DCLK1 also prohibits microtubule binding, implicating this domain in efficient lattice binding by DCLK1. Overall, our data lead to a model in which DCLK1 autophosphorylates its C-terminal tail to modulate the activity of its own kinase domain and subsequently the level of phosphorylation within its microtubule-binding domains. To our knowledge, this is the first example of a self-regulatory MAP that can tune its microtubule-binding properties based on autophosphorylation state. Our results uncover a novel intramolecular regulation of microtubule binding within a prominent family of MAPs and may have implications for DCLK1’s known roles in tumor development and cancer progression.

## Results

Previous results suggested that phosphorylation of DCLK1 occurs in part via autophosphorylation (37,39). To determine if DCLK1 phosphorylation is mediated by an inter-or intramolecular mechanism, we utilized an established kinase-dead mutant of DCLK1 (D511N)(39) and an active wild-type (WT) DCLK1 enzyme, both purified from bacteria (Figure S1B, S2A). We did not observe trans-phosphorylation of DCLK1-D511N upon incubation with DCLK1-WT, although DCLK1-WT efficiently autophosphorylated itself in this assay (Figure S2B). Thus, under the conditions in our experiments, DCLK1 phosphorylation occurs via an intramolecular mechanism.

Removal of the C-terminal region of DCLK1 that follows the kinase domain results in an increase in kinase activity (37). How this region regulates enzymatic activity and autophosphorylation of DCLK1, and how phosphorylation of the molecule affects its microtubule-binding properties are open questions. We first compared the mobility of full-length DCLK1-WT (aa 1-740) and a truncated DCLK1 lacking the C-terminal tail (ΔC: aa 1-648), to full-length kinase dead DCLK1-D511N (Figures 1A and S1B) on a Phos-tag gel, which enhances the separation of differentially phosphorylated proteins (41). We found that bacterially-expressed DCLK1-WT and DCLK1-ΔC proteins migrated more slowly into the Phos-tag gel, indicative of higher levels of phosphorylation, compared to the non-phosphorylated DCLK1-D511N (Figure 1A). Using total internal reflection fluorescence microscopy (TIRF-M), we imaged sfGFP-tagged WT, ΔC, and D511N proteins binding to taxol-stabilized microtubules (Figure 1B) at concentrations differing by 8-fold. Strikingly, DCLK1-ΔC did not bind to microtubules at either concentration tested, in stark contrast to DCLK1-WT and DCLK1-D511N, which both robustly bound to microtubules (Figure 1B-C). Notably, DCLK1-D511N bound microtubules more robustly at lower concentrations than DCLK1-WT. These experiments suggest that the C-terminal region regulates DCLK1 autophosphorylation, which in turn directly modulates its microtubule binding affinity.

**Figure 1.**
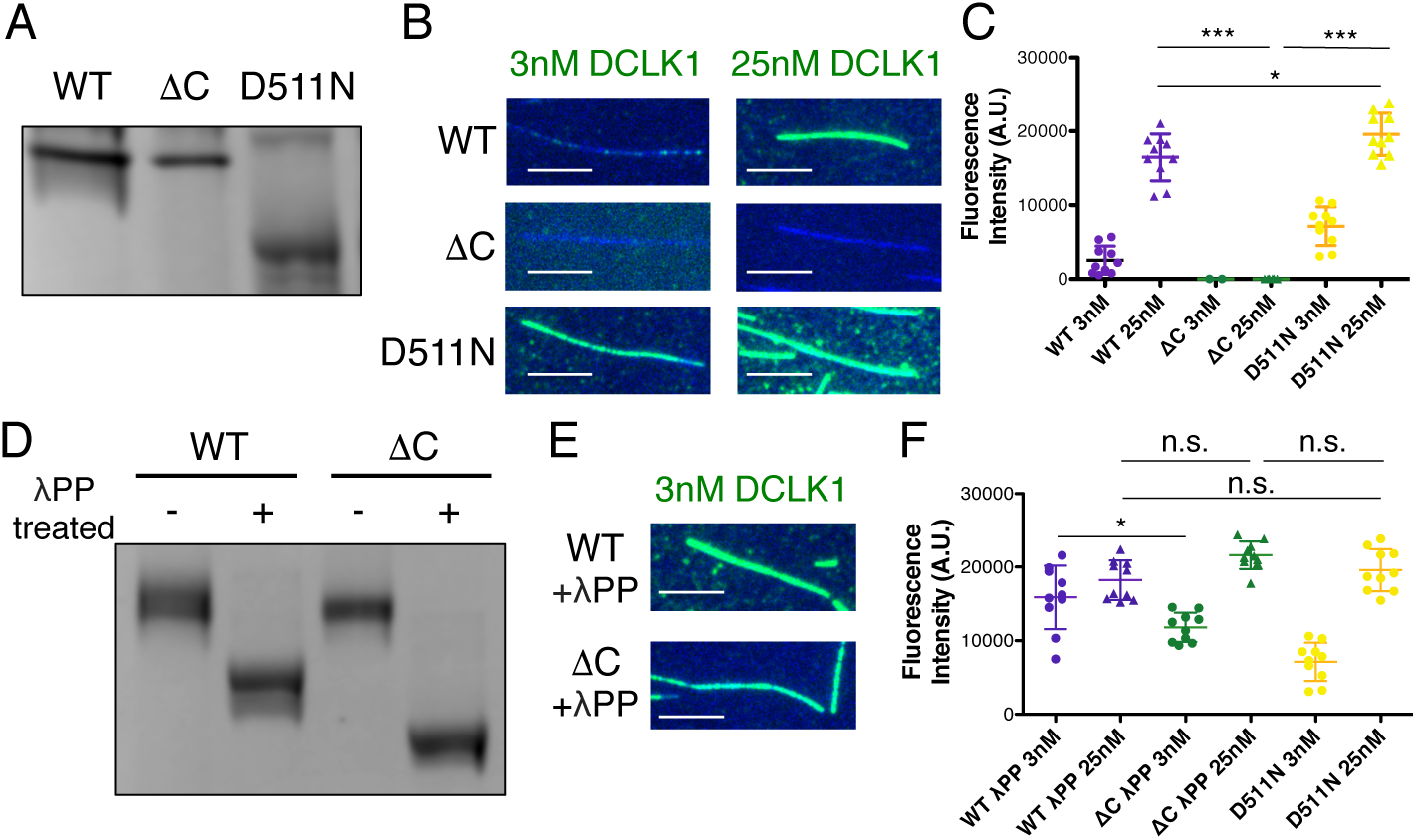
The C-terminal domain of DCLK1 regulates autophosphorylation and microtubule binding. (A) Coomassie Blue-stained SDS-PAGE Phos-tag gel of purified wild-type (WT), ΔC, and kinase dead (D511N) DCLK1 proteins separated by phosphorylation level. Representative gel from n = 3 independent experiments. (B) TIRF-M images of 3 nM and 25 nM sfGFP-DCLK1 WT, ΔC, and D511N (green), expressed in bacteria under standard conditions, binding to taxol-stabilized microtubules (blue). Scale bars: 5 μm. (C) Quantification of microtubule-bound sfGFP-DCLK1 fluorescence intensity. Means ± s.d.: 2555.9 ± 1902.3 for 3 nM WT, 16454.1 ± 3174.0 for 25 nM WT (n = 100 microtubules from n = 10 images from 2 independent trials for each concentration), 0.122 ± 8.3 for 3 nM ΔC, 3.82 ± 33.6 for 25 nM ΔC (n = 100 microtubules from n = 10 images from 2 independent trials for each concentration), 7137.4 ± 2611.8 for 3 nM D511N, and 19575.4 ± 2861.0 for 25 nM D511N (n = 100 microtubules from n = 10 images from 2 independent trials for each concentration). *** indicates *P* < 0.0001 and *P* = 0.0354 for 25 nM WT vs. 25 nM D511N, calculated by one-way ANOVA with Bonferroni correction. (D) Coomassie Blue-stained SDS-PAGE Phos-tag gel of purified DCLK1WT and ΔC incubated with lambda phosphatase (λPP) or incubated in buffer alone for 1 hr at 30°C. Representative gel from n = 2 independent experiments. (E) TIRF-M images of 3 nM sfGFP-DCLK1 WT and ΔC (green) after treatment with λPP, binding to taxol-stabilized microtubules (blue). Scale bars: 5 μm. (F) Quantification of microtubule-bound sfGFP-DCLK1 fluorescence intensity. Means ± s.d.: 15878.0 ± 4320.1 for 3 nM WT+λPP, 18233.2 ± 2683.9 for 25 nM WT+λPP (n = 100 microtubules from n = 10 images from 2 independent trials for each concentration), 11810.0 ± 1971.8 for 3 nM ΔC+λPP, and 21604.1 ± 1875.8 for 25 nM ΔC+λPP (n = 100 microtubules from n = 10 images from 2 independent trials for each concentration). D511N data are reproduced from Figure 1C for comparison. *P* = 0.1551 for 25 nM WT+λPP vs. 25 nM ΔC+λPP, *P* = 0.2935 for 25 nM WT+λPP vs. 25 nM D511N, *P* = 0.0771 for 25 nM ΔC+λPP vs.25 nM D511N, and *P* = 0.0339 for 3 nM WT+λPP vs. 3 nM ΔC+λPP, calculated by one-way ANOVA with Bonferroni correction.

To test this possibility, we sought to evaluate the microtubule binding behaviors of dephosphorylated DCLK1-WT and DCLK1-ΔC. We incubated the proteins with the Mn^2+^-dependent protein phosphatase, lambda phosphatase (λPP), which strongly dephosphorylated DCLK1-WT and DCLK1-ΔC as evidenced by marked shifts on a Phos-tag gel without phospho-intermediate bands (Figure 1D). The similar migration of DCLK1-WT and DCLK1-ΔC in the absence of phosphatase, coupled with the relatively larger migration shift of DCLK1-ΔC after treatment, further suggests that DCLK1-ΔC is hyperphosphorylated compared to the WT protein, in agreement with previous results (37). Using TIRF-M, we found that λPP-treated DCLK1-WT and DCLK1-ΔC bound to microtubules similarly to DCLK1-D511N (Figure 1E-F), further suggesting that autophosphorylation modulates the microtubule binding affinity of DCLK1 and that hyperphosphorylation of DCLK1-ΔC largely abolishes microtubule binding.

The high phosphorylation levels observed for both DCLK1-WT and DCLK1-ΔC suggested that these proteins phosphorylate themselves during bacterial expression. To determine the contributions of the C-terminal tail to DCLK1 function, we devised a strategy to control the levels of autophosphorylation during expression. We co-expressed DCLK1-WT and DCLK1-ΔC with λPP in bacteria, followed by subsequent removal of λPP from the DCLK1 preps via affinity and ion exchange chromatography. We compared DCLK1 proteins prepared in the absence or presence of λPP on a Phos-tag gel and observed that λPP co-expression substantially reduced phosphorylation levels of both DCLK1-WT and DCLK1-ΔC (Figure 2A). For all subsequent experiments, all DCLK1 protein variants were co-expressed with λPP. Upon incubation of dephosphorylated DCLK1 proteins with ATP, both DCLK1-WT and DCLK1-ΔC exhibited an increase in phosphorylation, but DCLK1-ΔC appeared to be entirely phosphorylated by 30 min, whereas DCLK1-WT displayed a number of phosphorylated intermediates even at 60 min (Figure 2A-B, 93.7 % of DCLK1-ΔC protein shifts to the uppermost band after a 30 min incubation with ATP compared with 41.8 % of DCLK1-WT protein after a 60 min incubation with ATP). Using TIRF-M, we determined the microtubule-binding affinities for DCLK1-WT and DCLK1-ΔC in the absence and presence of ATP (Figure 2C-E). We found that, in the absence of ATP, both proteins exhibited relatively similar microtubule-binding affinities (Figure 2C-E; 2.9 ± 0.5 nM vs. 3.9 ± 0.5 nM for WT and ΔC, respectively). After a 30 min incubation with ATP, the microtubule-binding affinity of DCLK1-WT moderately weakened, as evidenced by a 2-fold increase in K_D_ (Figure 2D; 6.0 ± 1.0 nM). However, incubation with ATP resulted in a dramatic ∼45-fold decrease in microtubule affinity of DCLK1-ΔC (Figure 2E; 174.3 ± 85.4 nM). These results indicate that the loss of its regulatory C-terminal tail results in aberrant DCLK1 hyperphosphorylation, leading to a dramatic loss of microtubule binding affinity. Thus, the kinase activity of DCLK1 directly controls its association with microtubules via intramolecular phosphorylation, which in turn is regulated by the C-terminus of the protein.

**Figure 2.**
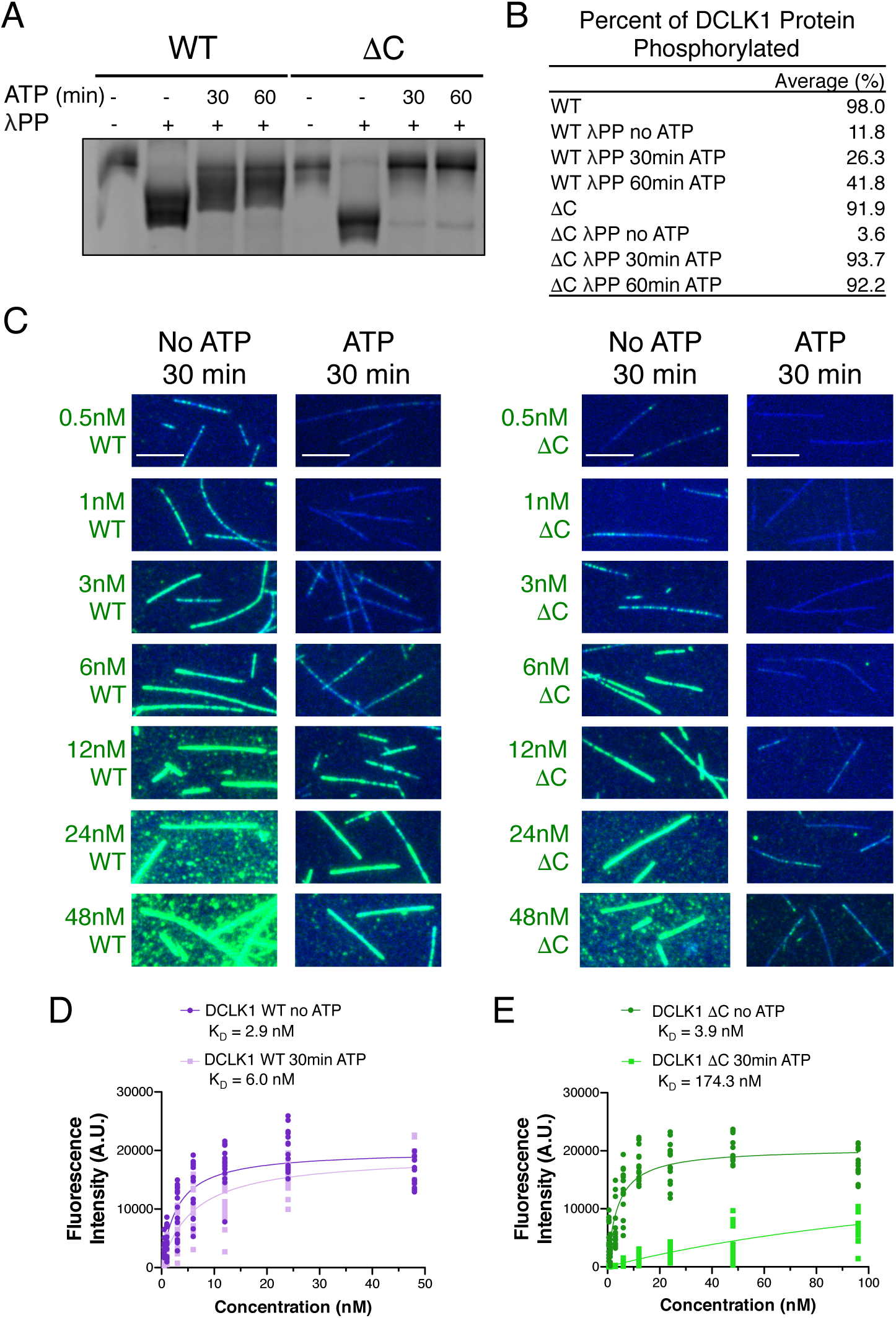
Hyperphosphorylation of DCLK1-ΔC prohibits microtubule binding. (A) Coomassie Blue-stained SDS-PAGE Phos-tag gel of purified DCLK1WT and ΔC proteins separated by phosphorylation level. The first and fifth lanes contain DCLK1 WT and ΔC expressed in bacteria under standard conditions. All other lanes contain DCLK1 WT and ΔC that were co-expressed with λPP, which was subsequently separated from DCLK1. Incubation of λPP-treated DCLK1 WT and ΔC with 2 mM ATP at the indicated times reveals a band shift indicative of an increase in phosphorylation. (B) Quantification of the average percent of total DCLK1 protein that is phosphorylated in each condition. Averages derived from n = 3 of independent experiments. (C) TIRF-M images of sfGFP-DCLK1 WT and ΔC (co-expressed in bacteria with λPP) at indicated concentrations (green) binding to taxol-stabilized microtubules (blue) after a 30 min incubation in the absence or presence of 2 mM ATP. Scale bars: 5 μm. (D) Quantification of microtubule-bound sfGFP-DCLK1-WT fluorescence intensity plotted against concentration after a 30 min incubation in the absence or presence of ATP (WT without ATP K_D_ ± s.d. = 2.9 ± 0.5 nM, n = 150 microtubules from n = 15 images from 3 independent trials for all concentrations; WT with ATP K_D_ ± s.d. = 6.0 ± 1.0 nM; n = 150, 150, 150, 150, 150, 150, and 100 microtubules for 0.5 nM, 1 nM, 3 nM, 6 nM, 12 nM, 24 nM, and 48 nM from n = 15 images from 3 independent trials; *P* < 0.0001 calculated using a student’s t-test). (E) Quantification of microtubule-bound sfGFP-DCLK1-ΔC fluorescence intensity plotted against concentration after a 30 min incubation in the absence or presence of ATP (ΔC without ATP K_D_ ± s.d. = 3.9 ± 0.5 nM; n = 150 microtubules from n = 15 images from 3 independent trials for each concentration; ΔC with ATP K_D_ ± s.d. = 174.3 ± 85.4 nM; n = 150 microtubules from n = 15 images from 3 independent trials for each concentration; *P* < 0.0001 calculated using a student’s t-test).

In order to determine how phosphorylation regulates the microtubule binding affinity of DCLK1-ΔC, we performed liquid chromatography with tandem mass spectrometry (LC-MS/MS) of phosphorylated DCLK1-WT and DCLK1-ΔC proteins. For each phosphorylated residue identified, we counted the total number of DCLK1-WT and DCLK1-ΔC peptides containing the phosphorylated residue, and then calculated the percent of those peptides whose spectra revealed phosphorylation at that residue. Within the microtubule-binding region of DCLK1 (aa 44-263), we found that 9 sites were more frequently phosphorylated in DCLK1-WT samples and 17 sites were more frequently phosphorylated in DCLK1-ΔC samples (Figure 3A-B). Of the 17 phosphorylation sites in DCLK1-ΔC, 5 either directly contact tubulin or are adjacent to residues that directly contact tubulin within the lattice (42). The architecture of DCLK1 suggests it likely has the flexibility to autophosphorylate its N-terminal half due to an intrinsically disordered region between the DC2 domain and the kinase domain (Figure 3C, aa 263-374). These results indicate that the loss of microtubule binding of DCLK1-ΔC is due to an increase in phosphorylation at multiple sites, as opposed to a single site whose phosphorylation status dictates microtubule binding.

**Figure 3.**
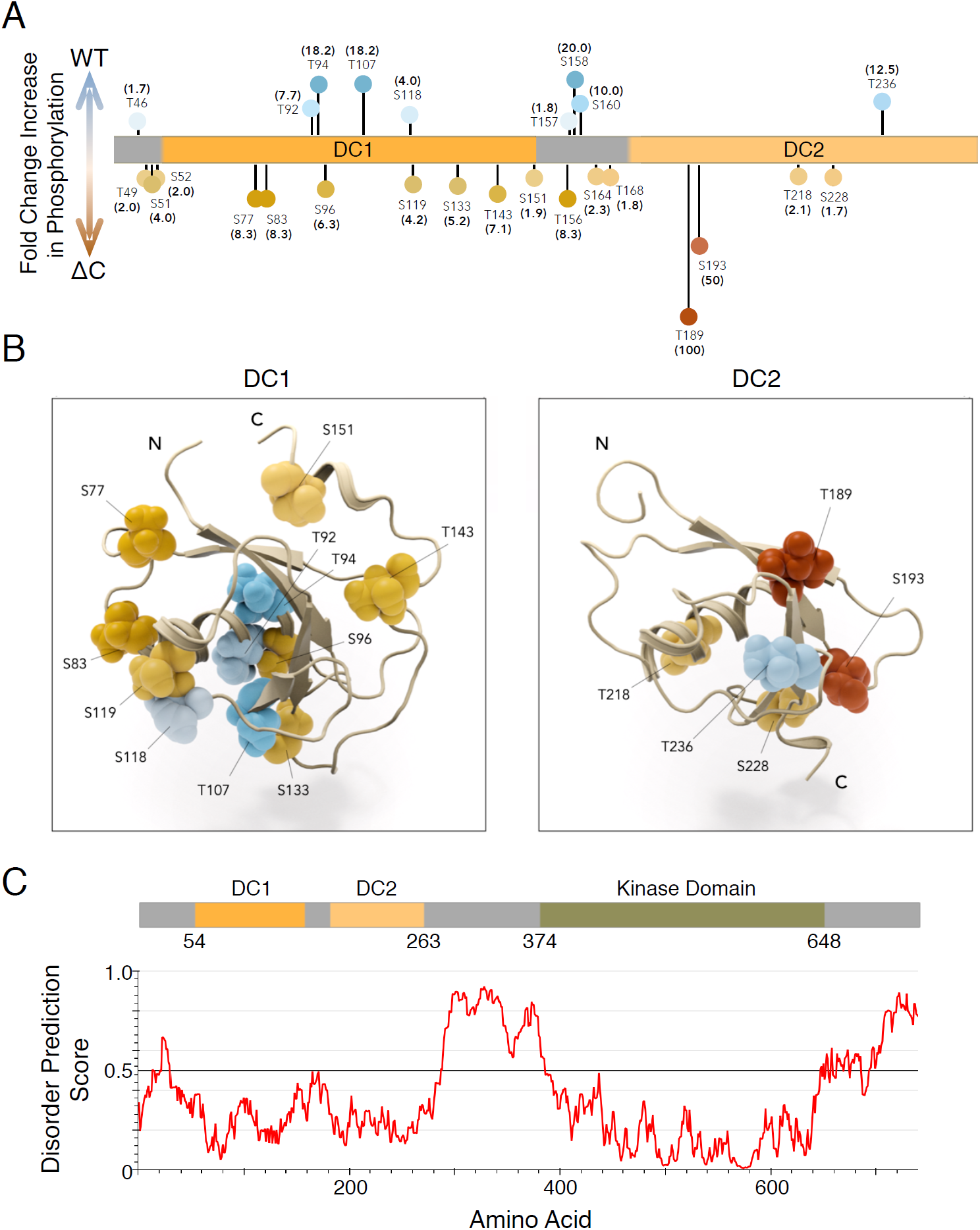
DCLK1-ΔC aberrantly autophosphorylates at multiple sites within the microtubule-binding region. (A-B) Visualization of changes in phosphorylation due to deletion of C-terminal domain. Experiment compared peptides from WT and ΔC constructs: data is expressed as fold change increases in phosphorylation in one construct over the other based on the percent of total peptides that exhibited phosphorylation at a particular site (n = 751 and 637 total peptides analyzed for WT and ΔC, respectively, from n = 3 independent experiments). (A) Lollipop plot summarizes changes in phosphorylation (≥ 1.5 fold change) mapped onto a diagram of DCLK1. (B) DC1 domain (1mg4, Kim et al. 2003) and DC2 domain modeled by homology to DCX-DC2 (5ip4, Burger et al. 2016 and Waterhouse et al. 2018). Domain structures are aligned and shown as ribbon representations with labeled S/T residues visualized as CPK/balls. Level of saturation in color indicates fold change in phosphorylation of those residues: increase in WT (blue colors) and increase in ΔC (orange/red colors). (C) Architecture of DCLK1 protein with the per-residue IUPRED2A {Meszaros, 2018 #3338} disorder prediction score shown in the corresponding plot with a cutoff value of 0.5 indicated by the dashed line. Residues scored above this value are predicted to be disordered.

Interestingly, the largest fold increase in DCLK1-ΔC phosphorylation occurred within the DC2 domain (Figure 3A-B). Prior cryo-EM work on the DCLK1 paralogue, DCX, has shown that only one DC domain binds stably to the microtubule lattice (30,43). Although originally hypothesized to be DC1, subsequent structural and biochemical studies called into question the identity of the bound DC domain (44,45). A recent cryo-EM study has proposed that DC2 binds to the newly polymerized GTP microtubule lattice, while DC1 binds the mature GDP microtubule lattice (42). All of our binding assays are performed with taxol-stabilized microtubules in the GDP state (46); however, based on our phosphorylation data, we reasoned that DC2 could still contribute to efficient binding of DCLK1 to GDP microtubules. In order to test if phosphorylation of the DC2 domain is responsible for the dramatic decrease in the microtubule-binding affinity of DCLK1-ΔC, we mutated 4 conserved residues within the DC2 domain DCLK1-ΔC_4A_) that showed an increase in phosphorylation and also sit at or adjacent to the tubulin-binding interface (Figure 4A, S3A)(42). We reasoned that if phosphorylation of these residues is responsible for the decreased microtubule-binding affinity of DCLK1-ΔC, then mutating these residues to alanines, which cannot be phosphorylated, should rescue the microtubule-binding defect of this construct in the presence of ATP. If, however, phosphorylation of these residues is not responsible for the decreased microtubule-binding affinity, then there should be no difference in binding between the DCLK1-ΔC and DCLK1-ΔC_4A_, regardless of the presence of ATP.

**Figure 4.**
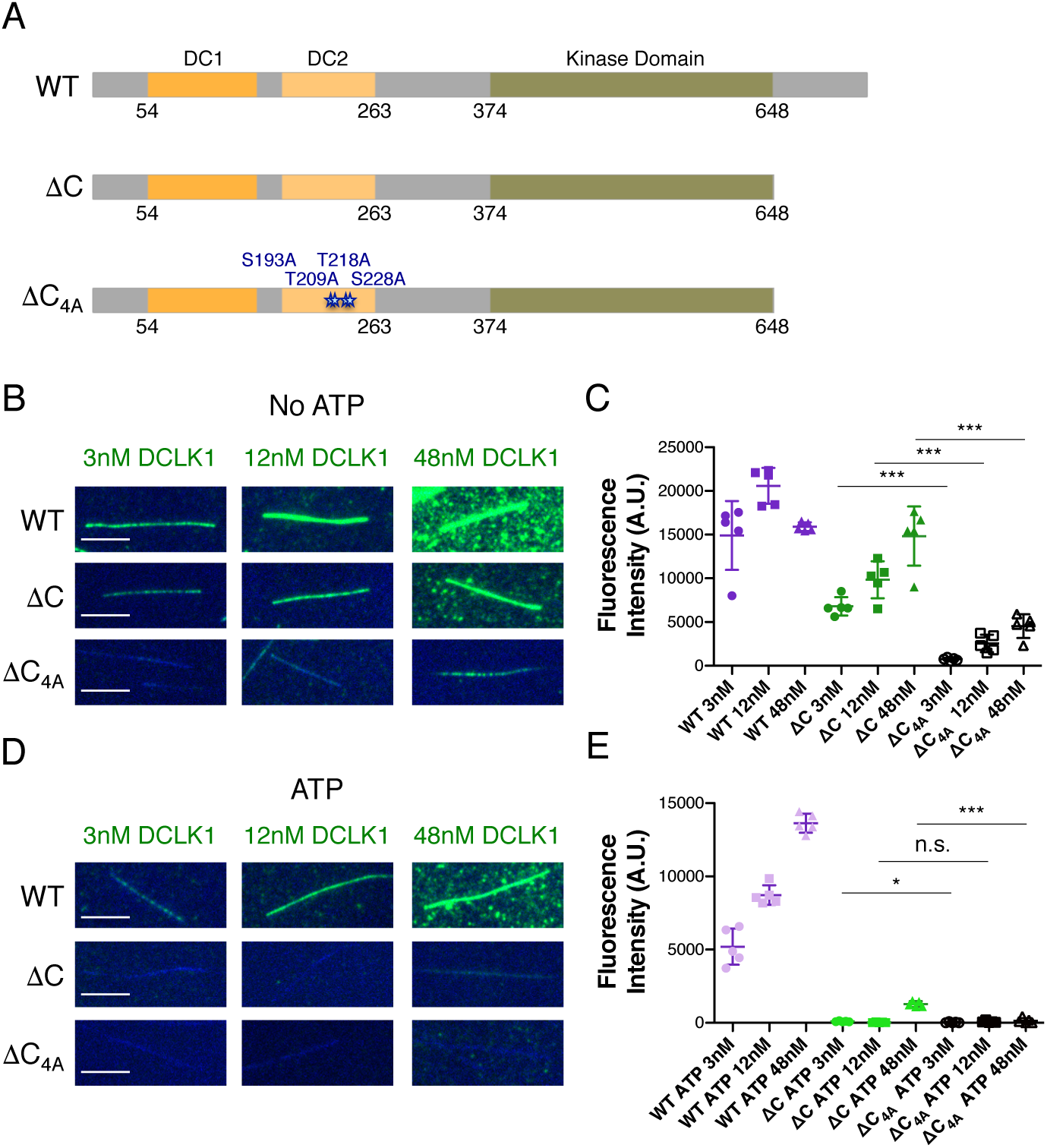
The DC2 domain of DCLK1 is necessary for microtubule binding. (A) Diagrams depicting the domains, amino acid boundaries, and mutations relevant to the DCLK1 constructs used for Figure 4. ΔC_4A_ indicates the four indicated residues in DC2 were mutated to alanines. Of these four residues, there was a ≥ 1.5 fold change in phosphorylation for S193, T218, and S228. Although the fold change for T209 was slightly under this threshold, it was still included in our mutational analysis due to its interface with the microtubule lattice. (B) TIRF-M images of sfGFP-DCLK1 WT, ΔC, and ΔC_4A_, co-expressed in bacteria with λPP, at indicated concentrations binding to taxol-stabilized microtubules (blue) in the absence of ATP. Scale bars: 5 μm. (C) Quantification of microtubule-bound sfGFP-DCLK1 fluorescence intensity. Means ± s.d.: 14908.5 ± 3935.8 for 3 nM WT, 20582.9 ± 2054.4 for 12 nM WT, 15905.6 ± 499.2 for 48 nM WT (n = 50 microtubules from n = 5 images for each concentration), 6792.6 ± 1049.4 for 3 nM ΔC, 9835.6 ± 2119.8 for 12 nM ΔC, 14827.6 ± 3384.0 for 48 nM ΔC (n = 50 microtubules from n = 5 images for each concentration), 744.7 ± 167.2 for 3 nM ΔC_4A_, 2547.3 ± 995.2 for 12 nM ΔC_4A_, 4529.5 ± 1349.6 for 48 nM ΔC_4A_ (n = 50 microtubules from n = 5 images for each concentation). *** indicates *P* < 0.0001 calculated by one-way ANOVA with Bonferroni correction. (D) TIRF-M images of sfGFP-DCLK1 WT, ΔC, and ΔC_4A_, co-expressed in bacteria with λPP then incubated for 30 min with 2mM ATP, at indicated concentrations binding to taxol-stabilized microtubules (blue). Scale bars: 5 μm. (E) Quantification of microtubule-bound sfGFP-DCLK1 fluorescence intensity. Means ± s.d.: 5195.3 ± 1230.4 for 3 nM WT+ATP, 8723.8 ± 668.9 for 12 nM WT+ATP, 13631.1 ± 648.2 for 48 nM WT+ATP (n = 50 microtubules from n = 5 images for each concentration), 95.3 ± 29.3 for 3 nM ΔC+ATP, 32.5 ± 26.5 for 12 nM ΔC+ATP, 1276.7 ± 200.2 for 48 nM ΔC+ATP (n = 50 microtubules from n = 5 images for each concentration), 28.0 ± 36.4 for 3 nM ΔC_4A_+ATP, 77.1 ± 72.2 for 12 nM ΔC_4A_+ATP, 142.1 ± 179.3 for 48 nM ΔC_4A_+ATP (n = 50 microtubules from n = 5 images for each concentration). *P* = 0.0122 for 3 nM ΔC+ATP vs. 3 nM ΔC_4A_+ATP; *P* = 0.2309 for 12 nM ΔC+ATP vs. 12 nM ΔC_4A_+ATP; *P* < 0.0001 for 48 nM ΔC+ATP vs. 48 nM ΔC_4A_+ATP, calculated by student’s t-test.

Using TIRF-M, we imaged increasing concentrations of DCLK-WT, DCLK1-ΔC, and DCLK1-ΔC_4A_ binding to taxol-stabilized microtubules in the presence or absence of ATP (Figure 4B-E). In contrast to DCLK1-ΔC, which binds robustly to microtubules in the absence of ATP, the binding of DCLK1-ΔC_4A_ was dramatically impaired in the absence of ATP (Figure 4B-C). This was a surprising result, because it indicates that the DC2 domain is indeed required for efficient binding not just to the GTP lattice, but also to the GDP lattice. In the presence of ATP, both DCLK1-ΔC and DCLK1-ΔC_4A_ showed a near total loss of microtubule affinity, consistent with our prior results in Figure 2C. Given the importance of these residues in interacting with the microtubule, these data also suggest that the addition of phosphate groups to these four residues would likely perturb microtubule binding. Therefore, aberrant phosphorylation of these residues in DCLK1-ΔC could indeed be responsible for the dramatic attenuation of microtubule binding.

We next wanted to determine the mechanism by which the C-terminal region of DCLK1 prevents hyperphosphorylation of the DC domains. We examined autophosphorylated DCLK1-WT by LC-MS/MS and identified two threonine residues in the C-terminal region (T687 and T688) that were consistently phosphorylated (Figure S3B). In order to understand how the C-terminal tail contributes to autophosphorylation, we mutated T687 and T688 to alanines individually and in combination (Figure 5A). For all of the experiments with these mutants, we co-expressed DCLK1 proteins with λPP to obtain a dephosphorylated protein preparation. We first evaluated the ability of these mutants to autophosphorylate using the Phos-tag gel system (Figure 5B-C). While DCLK1-WT and T687A exhibited a moderate increase in phosphorylation after a 30 min. incubation with ATP, T688A and T687/688A appeared to be entirely phosphorylated at this same time point (Figure 5B-C, 13.2 %, 62.7 %, 96.0 %, and 90.1 % of protein shifts to the most phosphorylated band after a 30 min incubation with ATP for WT, T687A, T688A, and T687/688A, respectively). Therefore, phosphorylation of T688 within the C-terminal domain may be critical for the regulation of the kinase activity of DCLK1. To further elucidate the consequences of abolishing these phosphorylation sites, we used TIRF-M to determine microtubule-binding affinities for the DCLK1 phospho-mutants in the presence or absence of ATP (Figure 5D-F). In the absence of ATP, all DCLK1 proteins, with the exception of T687/688A double mutant, exhibited relatively similar microtubule-binding affinities based on the dissociation constants derived from fluorescent saturation curves (Figure 5D-F; K_D_ = 2.9 ± 0.4 nM, 2.4 ± 0.4 nM, 2.4 ± 0.4 nM, 9.4 ± 1.4 nM for WT, T687A, T688A, and T687/688A, respectively). After a 30 min. incubation with ATP, the microtubule-binding affinities of DCLK1-WT and T687A were comparable, whereas T688A and T687/688A displayed a dramatic reduction in microtubule binding with a ∼10-fold and ∼36-fold increase in K_D_, respectively (Figure 5F; 25.7 ± 7.5 nM for T688A and 341.1 ± 565.3 nM for T687/688A). These results indicate that DCLK1 likely autophosphorylates residues within its C-terminal region in order to control aberrant hyper-phosphorylation within its microtubule-binding domain.

**Figure 5.**
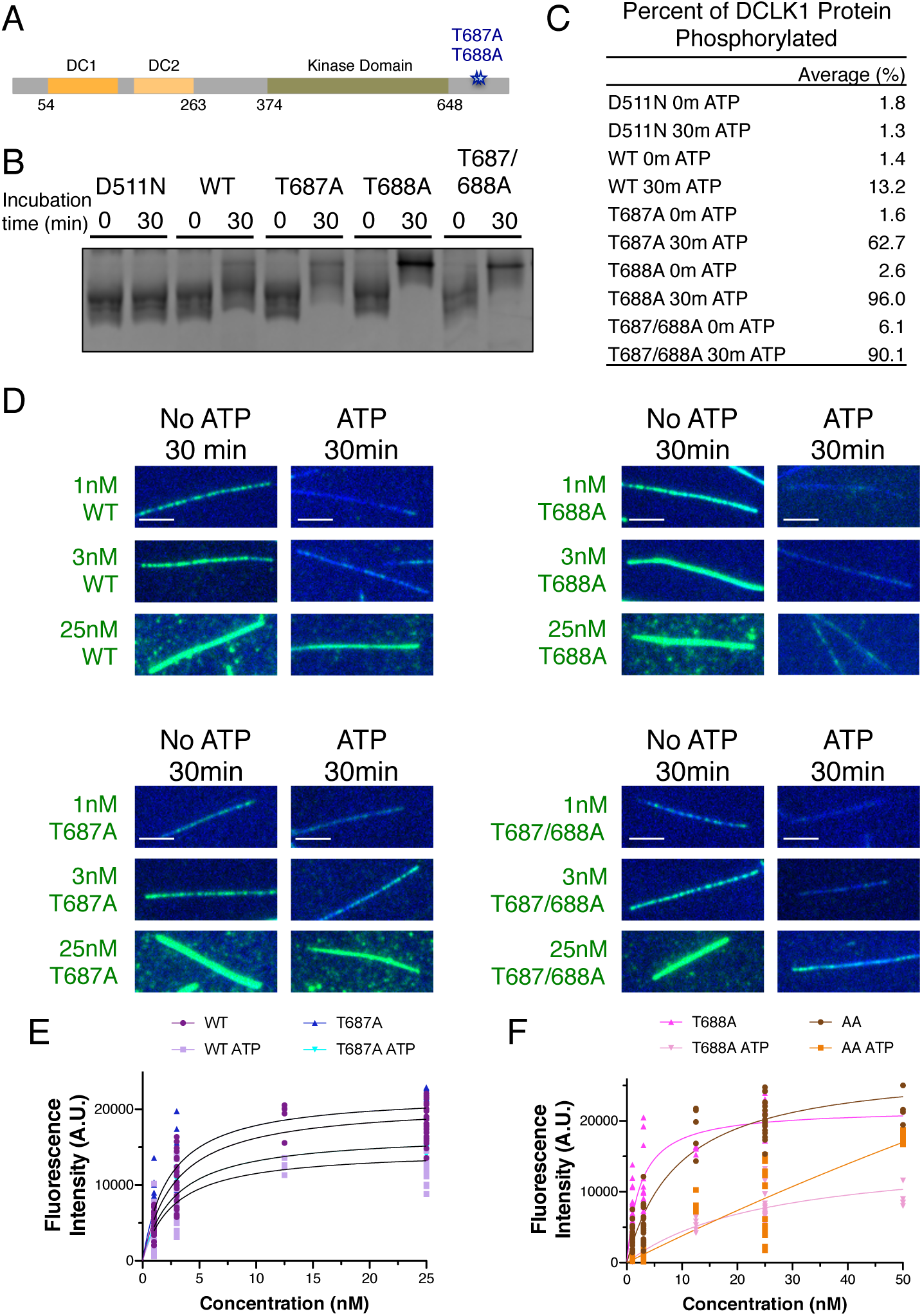
Normal autophosphorylation within the C-terminal domain of DCLK1 is necessary to prevent hyperphosphorylation of the rest of the molecule. (A) Diagram depicting the domains, amino acid boundaries, and mutations in the C-terminal region relevant to the DCLK1 constructs used for Figure 5. (B) Coomassie Blue-stained SDS-PAGE Phos-tag gel of purified kinase dead (D511N), WT, T687A, T688A, and T687/688A DCLK1 proteins separated by phosphorylation level. For all experiments, DCLK1 proteins were co-expressed with λPP, which was subsequently separated from DCLK1. Incubation of λPP-treated DCLK1 proteins with 2 mM ATP at the indicated times reveals a shift in the phosphorylation level to varying degrees. (C) Quantification of the average percent of total DCLK1 protein that is phosphorylated in each condition. Averages derived from n = 3 of independent experiments. (D) TIRF-M images of sfGFP-DCLK1 WT, T687A, T688A, and T687/688A (co-expressed in bacteria with λPP) at indicated concentrations (green) binding to taxol-stabilized microtubules (blue) after a 30 min. incubation in the absence or presence of 2 mM ATP. Scale bars: 5 μm. (E) Quantification of microtubule-bound sfGFP-DCLK1 WT and T687A fluorescence intensity plotted against concentration after a 30 min incubation in the absence or presence of ATP (WT without ATP K_D_ ± s.d. = 2.9 ± 0.4 nM; n = 200, 200, 50, and 200 microtubules for 1 nM, 3 nM, 12.5 nM, and 25 nM concentrations, respectively, from 4 independent trials; WT with ATP K_D_ ± s.d. = 2.7 ± 0.5 nM; n = 200, 200, 50, and 200 microtubules for 1 nM, 3 nM, 12.5 nM, and 25 nM concentrations, respectively, from 4 independent trials; T687A without ATP K_D_ ± s.d. = 2.4 ± 0.4 nM; n = 150, 150, and 150 microtubules for 1 nM, 3 nM, and 25 nM concentrations, respectively from 3 independent trials; T687A with ATP K_D_ ± s.d. = 2.8 ± 0.4 nM; n = 150, 150, and 150 microtubules for 1 nM, 3 nM, and 25 nM, from 3 independent trials; *P* = 0.7885 for WT+ATP vs. T687A+ATP using a student’s t-test). (F) Quantification of microtubule-bound sfGFP-DCLK1 T688A and T687/688A fluorescence intensity plotted against concentration after a 30 min. incubation in the absence or presence of ATP (T688A without ATP K_D_ ± s.d. = 2.4 ± 0.4 nM; n = 200, 200, 50, 200, and 50 microtubules for 1 nM, 3 nM, 12.5 nM, 25 nM, and 50 nM concentrations, respectively from 4 independent trials; T688A with ATP K_D_ ± s.d. = 25.7 ± 7.5 nM; n = 200, 200, 50, 190, and 50 microtubules for 1 nM, 3 nM, 12.5 nM, 25 nM, and 50 nM concentrations, respectively, from 4 independent trials; T687/688A without ATP K_D_ ± s.d. = 9.4 ± 1.4 nM; n = 200, 200, 50, 200, and 50 microtubules for 1 nM, 3 nM, 12.5 nM, 25 nM, and 50 nM concentrations, respectively from 4 independent trials; T687/688A with ATP K_D_ ± s.d. = 341.4 ± 565.3 nM; n = 200, 200, 50, 200, and 50 microtubules for 1 nM, 3 nM, 12.5 nM, 25 nM, and 50 nM concentrations, respectively, from 4 independent trials; *P* = 0.0009 for WT+ATP vs. T688A+ATP and *P* = 0.2760 for WT+ATP vs. T687/688A+ATP using a student’s t-test).

There could be consequences of autophosphorylation for DCLK1 function other than microtubule binding. We therefore analyzed the effect of autophosphorylation on DCLK1 conformation and on DCLK1 sensitivity to calpain cleavage. First, we fractionated the following proteins by sucrose density centrifugation: normally expressed DCLK1-WT, DCLK1-WT co-expressed with λPP, DCLK1-D511N, and DCLK1-T688A co-expressed with λPP (Figure S4A-B). We observed a similar profile for all proteins on a 3-9 % step gradient (Figure S4A-B), indicating these proteins adopt similar conformations regardless of phosphorylation state. DCLK1 contains two PEST domains that are targeted by calpain for proteolytic cleavage (47). Upon incubation of calpain with DCLK1-WT expressed either under normal conditions or co-expressed with λPP, we detected similar cleavage products of DCLK1 under both conditions (Figure S4C). Therefore, autophosphorylation does not affect cleavage of purified DCLK1-WT by calpain under our conditions. These results support a regulatory role for autophosphorylation in dictating the microtubule-binding affinity of DCLK1, without affecting the overall conformation of the molecule or its sensitivity to calpain cleavage.

## Discussion

Overall, our study elucidates a mechanism by which DCLK1 modulates its kinase activity to tune its microtubule-binding affinity. We have found that DCLK1 autophosphorylates within its C-terminal tail to prevent aberrant hyper-phosphorylation within its microtubule-binding domain. To our knowledge, this is the first example of a MAP whose binding is controlled by autophosphorylation. Based on the relevancy of DCLK1 to the progression of a variety of cancer types, understanding DCLK1 autoregulation is critical in determining its biological function in healthy versus disease states.

Autophosphorylation control of microtubule-binding affinity is likely to be controlled by cellular context. DCX is tightly regulated at the transcriptional level to ensure proper temporal expression during neuronal development (22,38,48,49). In contrast, members of the DCLK family are expressed during embryonic, post-embryonic, and adult periods, and are also expressed in a range of tissues outside the brain (18,38). This lack of temporal restriction may therefore necessitate auto-regulation, as well as other modes of control, for the DCLK family in a range of cellular activities.

Although DCX and DCLKs have been found to nucleate, tip-track, and bundle microtubules, the physiological functions of these proteins in regulating microtubule growth and organization remain elusive (30-33,36). Understanding the conserved mechanisms by which these paralogues bind to microtubules is essential in ascribing molecular functions to DCX family members, most of which are implicated in disease (18). Prior work has demonstrated that isolated DC domains cannot stimulate microtubule polymerization or effectively bind microtubules, indicating both domains are required in tandem (23-25,50). The contributions of the individual DC domains are still controversial. Initial cryo-EM structures of DCX on microtubules revealed only a single bound DC domain, which was hypothesized to be DC1 (30,43). However, a subsequent study showed that specifically blocking DC2, but not DC1, prevented DCX from interacting with microtubules, suggesting DC2 is critical for lattice binding (44). Recent cryo-EM data of DC1 and DC2 bound to microtubule lattices in different nucleotide states unveiled that DC2 binds to the GTP microtubule lattice, while DC1 prefers the GDP microtubule lattice (42). This model is consistent with observations that DCX tracks the growing plus-end of the microtubule, which is specifically disrupted by missense mutations in DC2 (33). It was therefore surprising that mutating 4 residues in DC2 abolished DCLK1 binding to GDP-taxol microtubules (Figure 4). While DC2 may preferentially recognize the GTP state of polymerized tubulin, our data indicate that it may also be important for the molecule to stably interact with the GDP lattice. We hypothesize that DC2 could be necessary for initial interactions with the lattice, regardless of nucleotide state, or for the cooperative binding exhibited by members of the DCX family (32). It will be imperative in the future to determine the individual and tandem roles of the DC domains across the DCX superfamily in a range of microtubule processes.

We have found that phosphorylation at different sites within DCLK1 have distinct effects on the molecule. Phosphorylation within the C-terminal region is essential to restrict the kinase activity of DCLK1, preventing hyper-phosphorylation within the DC2 domain and eradication of microtubule binding. This mechanism could prove important in directing the kinase activity of DCLK1 to orthogonal molecular substrates (51) instead of itself. Alternatively, the C-terminal tail of DCLK1 could be cleaved under specific cellular situations to release it from the microtubule. Our assays provide evidence that autophosphorylation does not appear to affect calpain cleavage or the overall conformation of DCLK1. However, these results do not rule out effects of phosphorylation on processing by other proteolytic enzymes, or potentially rapid, dynamic conformational changes that would not be apparent in our assays. Future studies will be necessary to expand upon these results both in vitro and in vivo and determine how DCLK1 function is regulated within the cell.

The region of DCLK1 spanning the kinase domain and C-terminal tail is 46% identical to the comparable region of CamKI (37-39). The flexible C-terminal tail of CamKI serves as a regulatory switch; it forms multiple interactions with the kinase domain and keeps it in an inactive conformation (40). Our data implicating the C-terminal tail of DCLK1 in preventing kinase hyperactivity combined with prior structural data propose a similar model for DCLK1 (39). Furthermore, we have identified a residue (T688) within the DCLK1 C-terminal tail that is critical in modulating kinase activity. Whether the DCLK1 C-terminal tail interacts stably or dynamically with the kinase domain, and when DCLK1 may need to switch from a microtubule bound to unbound state within the cell are open questions. Overall, the evidence for a conserved mechanism governing kinase activity for CaMKI and DCLK1 and how this could go awry in disease provide exciting new avenues for future exploration.

Finally, this study has implications for one of the greatest human adversaries: cancer. There are over 100 discrete forms of cancer, each with multiple causes (52). Numerous studies have found that DCLK1 is upregulated and acts as an oncogene in a range of cancers including pancreatic, colorectal, gastric, bladder, and breast cancer (7-14). Due to the emerging body of evidence implicating DCLK1 in tumorigenesis, the protein appears to be a promising target for not just one, but for several types of cancers (15-17). However, the complex intramolecular mechanism of DCLK1 must be thoroughly dissected before the field will be able to develop therapeutically effective drugs. For example, in light of our work, developing kinase inhibitors may not prove to be an effective means of controlling DCLK1’s microtubule-binding functions, because WT and kinase dead DCLK1 bind with similar affinities to microtubules. Future studies on the biological functions of DCLK1 microtubule binding and kinase activity during the initiation and progression of cancer cell proliferation and migration will provide fundamental insights into how DCLK1 contributes to this malady and how it can be adequately targeted.

## Experimental Procedures

### Molecular Biology

The cDNAs for protein expression in this study were as follows: DCLK1 (Transomic, BC133685) and Lambda phosphatase (Addgene, 79748). DCLK1 proteins were cloned in frame using Gibson cloning into a pET28 vector with an N-terminal strepII-Tag and a superfolder GFP (sfGFP) cassette. Lambda phosphatase protein was cloned in frame using Gibson cloning into a pET11 vector with an N-terminal RFP cassette.

### Protein Expression and Purification

Tubulin was isolated from porcine brain using the high-molarity PIPES procedure as previously described (53). For bacterial expression of all sfGFP-DCLK1 variants, BL21 cells were grown at 37°C until ∼O.D. 0.6 and protein expression was induced with 0.1 mM IPTG. For DCLK1 variants co-expressed with RFP-lambda phosphatase, BL21 cells were co-transformed with sfGFP-DCLK1 and RFP-lambda phosphatase and grown using the same protocol. Cells were grown overnight at 18°C, harvested, and frozen. Cell pellets were resuspended in lysis buffer (50 mM Tris pH 8.0, 150 mM K-acetate, 2 mM Mg-acetate, 1 mM EGTA, 10% glycerol) with protease inhibitor cocktail (Roche), 1 mM DTT, 1 mM PMSF, and DNAseI. Cells were then passed through an Emulsiflex press and cleared by centrifugation at 23,000 x g for 20 mins. Clarified lysate from bacterial expression was passed over a column with Streptactin XT Superflow resin (Qiagen). After incubation, the column was washed with four column volumes of lysis buffer, then bound proteins were eluted with 50 mM D-biotin (CHEM-IMPEX) in lysis buffer (pH 8.5). Eluted proteins were concentrated on Amicon concentrators and passed through a HiTrap Q HP anion exchange chromatography column in lysis buffer using a Bio-Rad NGC system. Peak fractions were collected, concentrated, and flash frozen in LN_2_. Protein concentration was determined by measuring the absorbance of the fluorescent protein tag and calculated using the molar extinction coefficient of the tag. The resulting preparations were analyzed by SDS polyacrylamide gel electrophoresis (SDS-PAGE).

### Autophosphorylation Assays

For autophosphorylation assays to determine an intra-vs. inter-molecular mechanism, 500 nM of WT and/or kinase dead (D511N) DCLK1 proteins were incubated in the absence or presence of 0.5 mM ATPγS (Fisher 11912025MG) in assay buffer containing 50 mM Tris pH 8.5, 50 mM K-acetate, 2 mM Mg-acetate, 1 mM EGTA, and 10% glycerol, supplemented with 1 mM DTT and 1 mM PMSF for 30 min at 37°C. To determine if DCLK1-WT trans phosphorylates DCLK1-D511N, we cleaved the strepII-sfGFP tag off of the DCLK1-WT protein using TEV protease and subsequently subjected the DCLK1-WT to gel filtration to separate it from the protease. All samples were quenched with 0.1 mM EDTA, then incubated at room temperature with 2.5 mM p-Nitrobenzyl mesylate (Abcam ab138910) for ∼1 hr. All samples were then run on an SDS-PAGE gel, transferred to a PVDF membrane using an iBlot2 at 25V for 7 min. The membrane was immunoblotted with primary antibodies: mouse anti-strep (1:2500, Fisher NBP243719) and rabbit anti-thiophosphate ester (1:2000, Abcam ab133473), washed, incubated with secondary antibodies: Alexa 680 goat anti-mouse (1:10,000, Fisher A28183) and Dylight 800 goat anti-rabbit (1:10,000, Rockland labs 611-145-002), washed, then imaged on a LiCor Odyssey.

For autophosphorylation assays using the Phos-tag gel system, assays were performed using adenosine triphosphate (ATP) in assay buffer containing 50mM Tris, pH 8.5, 50mM K-acetate, 2mM Mg-acetate, 1mM EGTA, and 10% glycerol, supplemented with 1mM DTT and 1mM PMSF. The samples were incubated at room temperature for 30 or 60 minutes in the presence of assay buffer with 500nM DCLK1 and 2mM ATP. Non-phosphorylated samples included assay buffer and 500nM DCLK1. All samples were run on a Phos-Tag gel and analyzed as described below.

### Phos-Tag Gel Assays

Purified proteins were separated using Phos-tag gel technology (Wako, Phos-tag AAL-107). Samples were either expressed alone in BL21 cells and purified, treated with lambda phosphatase protein after purification at 30°C for 1 hr (New England BioLabs, P0753L), or co-expressed in BL21 cells with RFP-lambda phosphatase. Purified protein samples were incubated in the presence or absence of ATP in kinase assay protein buffer (50 mM Tris pH 8.5, 50 mM K-acetate, 2 mM Mg-acetate, 1 mM EGTA, 10% glycerol) with 1 mM DTT and 1 mM PMSF for 30 or 60 minutes at room temperature. Phos-tag SDS-PAGE was performed with pre-cast 7.5% polyacrylamide gels containing 50 μM Phos-tag acrylamide with MnCl_2_ (Wako, 192-18001). Electrophoresis was completed at 180 v for 90 min and the gel was stained with Coomassie Blue. The stained gel was imaged using a GelDoc (BioRad) and band intensity was quantified using ImageJ to draw a box over both the highest band and the lowest band in each lane. The measure for percent of protein that was phosphorylated was generated by dividing the intensity value of the high band by the total intensity from the sum of the high and low band.

### TIRF Microscopy

For TIRF-M experiments, a mixture of native tubulin, biotin-tubulin, and fluorescent-tubulin purified from porcine brain (∼10:1:1 ratio) was assembled in BRB80 buffer (80mM PIPES, 1mM MgCl_2_, 1mM EGTA, pH 6.8 with KOH) with 1mM GTP for 15 min at 37°C, then polymerized MTs were stabilized with 20 μM taxol. Microtubules were pelleted over a 25% sucrose cushion in BRB80 buffer to remove unpolymerized tubulin. Flow chambers containing immobilized microtubules were assembled as described (54). Imaging was performed on a Nikon Eclipse TE200-E microscope equipped with and Andor iXon EM CCD camera, a 100X, 1.49 NA objective, four laser lines (405, 491, 568, and 647 nm) and Micro-Manager software (55). All experiments were performed in assay buffer (60mM Hepes pH 7.4, 50mM K-acetate, 2mM Mg-acetate, 1mM EGTA, and 10% glycerol) supplemented with 0.1mg/mL biotin-BSA, 0.5% Pluronic F-168, and 0.2mg/mL κ-casein (Sigma) and 10μM taxol.

For all saturation curves, a concentration series was performed for each protein. For fluorescence intensity analysis, ImageJ was used to draw a line across the microtubule of the DCLK1 channel and the integrated density was measured. The line was then moved adjacent to the microtubule of interest and the local background was recorded. The background value was then subtracted from the value of interest to give a corrected intensity measurement. The fluorescence intensity data were fit with a one site binding hyperbola equation to derive the K_D_ for each DCLK1 variant.

### Mass Spectrometry

Samples were prepared for mass spectrometry analysis by incubating each DCLK1 variant with ATP for time periods ranging from 5 to 15 minutes. The reaction was quenched with 10mM EDTA. Protein of interest was first reduced at 56ºC for 45 minutes in 5.5 mM DTT followed by alkylation for one hour in the dark with iodoacetamide added to a final concentration of 10 mM. Trypsin was added at a final enzyme:substrate mass ratio of 1:50 and digestion carried out overnight at 37ºC. The reaction was quenched by flash freezing in liquid nitrogen and the digest was lyophilized. Digest was reconstituted in 0.1% TFA with 10% acetonitrile prior to injection.

The mass spectrometry instrument used to analyze the samples was a Xevo G2 QTof coupled to a nanoAcquity UPLC system (Waters, Milford, MA). Samples were loaded onto a C18 Waters Trizaic nanotile of 85 um × 100 mm; 1.7 µm (Waters, Milford, MA). The column temperature was set to 45°C with a flow rate of 0.45 mL/min. The mobile phase consisted of A (water containing 0.1% formic acid) and B (acetonitrile containing 0.1% formic acid). A linear gradient elution program was used: 0–40 min, 3– 40 % (B); 40-42 min, 40–85 % (B); 42-46 min, 85 % (B); 46-48 min, 85-3 % (B); 48-60 min, 3% (B).

Mass spectrometry data were recorded for 60 minutes for each run and controlled by MassLynx 4.1 (Waters, Milford, MA). Acquisition mode was set to positive polarity under resolution mode. Mass range was set form 50 – 2000 Da. Capillary voltage was 3.5 kV, sampling cone at 25 V, and extraction cone at 2.5 V. Source temperature was held at 110C. Cone gas was set to 25 L/h, nano flow gas at 0.10 Bar, and desolvation gas at 1200 L/h. Leucine–enkephalin at 720 pmol/ul (Waters, Milford, MA) was used as the lock mass ion at m/z 556.2771 and introduced at 1 uL/min at 45 second intervals with a 3 scan average and mass window of +/-0.5 Da. The MS^e^ data were acquired using two scan functions corresponding to low energy for function 1 and high energy for function 2. Function 1 had collision energy at 6 V and function 2 had a collision energy ramp of 18−42 V.

RAW MS^e^ files were processed using Protein Lynx Global Server (PLGS) version 2.5.3 (Waters, Milford, MA). Processing parameters consisted of a low energy threshold set at 200.0 counts, an elevated energy threshold set at 25.0 counts, and an intensity threshold set at 1500 counts. The databank used was derived from human. Searches were performed with trypsin specificity and allowed for two missed cleavages. Possible structure modifications included for consideration were methionine oxidation, carbamidomethylation of cysteine, and phosphorylation of serine, threonine, or tyrosine.

For viewing, PLGS search results were exported in Scaffold v4.4.6 (Proteome Software Inc., Portland, OR)

### Sucrose Gradients

Three-step sucrose gradients were prepared in centrifuge tubes using 250μL steps of 3%, 6%, and 9% sucrose in protein buffer and allowed to sit overnight at 4°C. The next morning, 100μL of ∼600nM of each DCLK1 variant was layered on top of the sucrose gradient, which were centrifuged at 50,000rpm for 4 hours at 4°C using a TLS55 rotor and an Optima MAX-XP Ultracentrifuge (Beckman Coulter). The sample was physically fractionated into 10 fractions of 85 uL each, which were run on an SDS-PAGE gel to assess protein sedimentation location. For quantification of protein levels in each fraction, ImageJ was used to draw a box of consistent size over the band in each lane and total intensity within the box was measured. The intensity of the band in each lane was divided by the total intensity of all boxes combined to produce the percent of protein in each fraction.

### Calpain Cleavage Assay

Calpain cleavage assays were performed in assay buffer containing 20 mM HEPES pH 7.2, 100 mM KCl, and 2 mM CaCl_2_. The samples were incubated at 30°C for 15 or 30 minutes in the presence of 750 nM DCLK1 and 0.617 μg human calpain 1 (Sigma, C6108), then analyzed by SDS-PAGE.

### Statistical Analysis

Statistical tests were performed with a two-tailed unpaired Student’s t-test, or a one-way ANOVA with Bonferroni post-hoc correction. Unless otherwise stated, all data was analyzed manually using ImageJ (FIJI). Graphs were created using Graphpad Prism and statistical tests were performed using this program. All variances given represent standard deviation. The statistical details of each experiment can be found in the figure legends.

### Data Availability

The data that support the findings of this study are available from the corresponding author upon reasonable request.

## Acknowledgements

We thank Richard McKenney and members of the Ori-McKenney and McKenney labs for reading the manuscript and providing feedback.

## Author Contributions

M.R., A.R., and K.M.O.M. conceived of the project and designed the experiments. M.R., A.R., A.D., J.C. and H.B. cloned all DCLK1 constructs and purified the recombinant proteins. M.R. performed the in vitro TIRF-M experiments, the kinase assays, the sucrose gradients, the cleavage assays, and analyzed all of data. A.R. performed the ATPγS kinase assays. D.W.N. created the molecular models. M.R. and K.M.O.M. wrote the manuscript with input from all authors.

## Supporting Information

**Figure S1.**
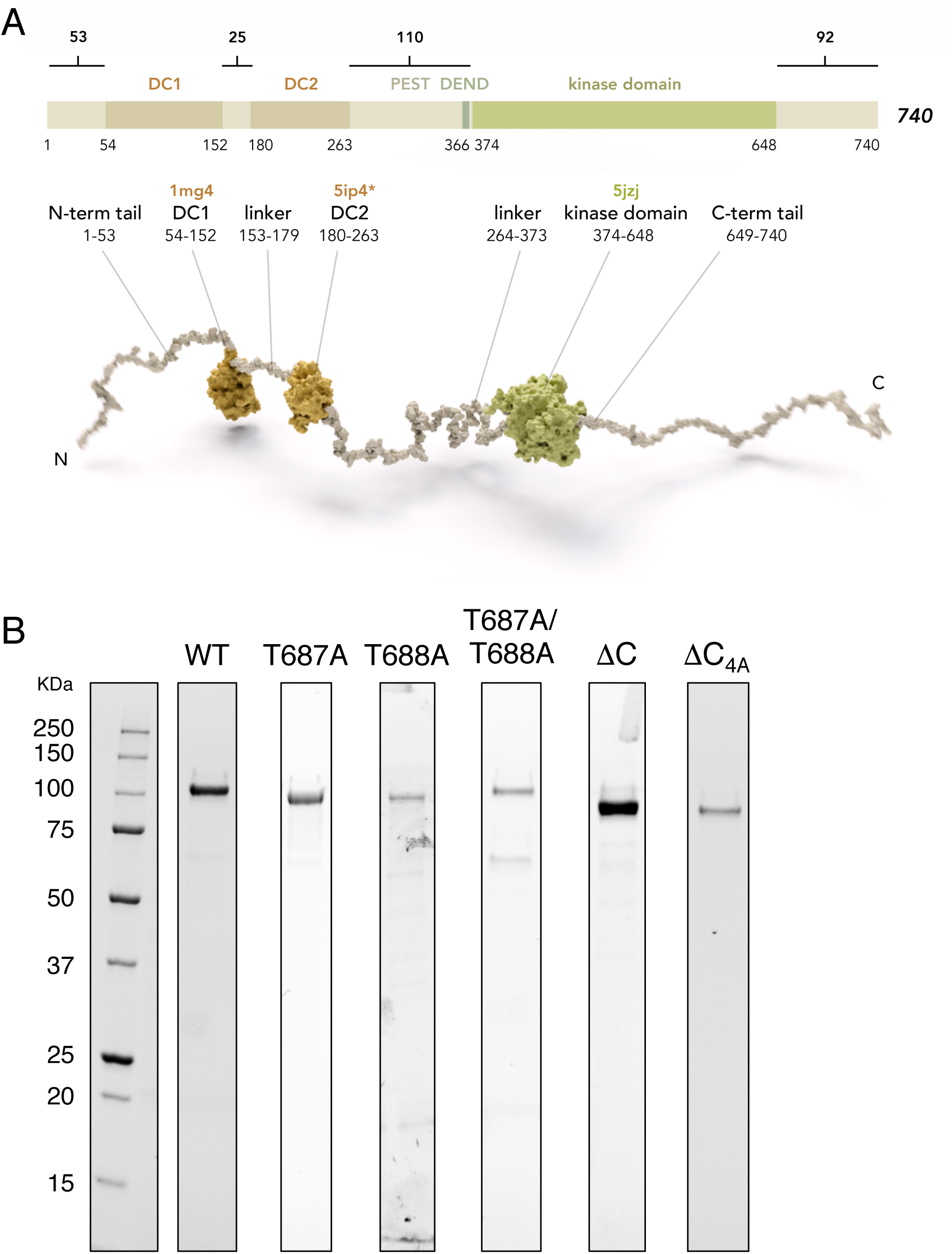
Diagram of full-length DCLK1 used in this study and gels for purified recombinant proteins used in this study. Related to Figures 1-5. (A) Diagram of domains and motifs of human DCLK1 (UniProt O15075). DC1, N-terminal Doublecortin-like (DCX) domain; DC2, C-terminal DCX domain; kinase domain. Motifs enriched in PEST (proline/P, glutamic acid/E, serine/S, threonine/T); DEND (aspartic acid/D, glutamic acid/E, asparagine/N, aspartic acid/D) based on Burgess et al. (2001) and Nagamine et al. (2011). Below: model of human DCLK1. DC1 domain (1mg4, Kim et al. 2003), DC2 domain modeled by homology to DCX-DC2 (5ip4, Burger et al. 2016 and Waterhouse et al. 2018), kinase domain (5jzj, Patel et al. 2016). DCLK1 is shown as a full-length pseudo-model, with projection domains/tails (and domain linkers) modeled as unfolded to visualize length and convey intrinsic disorder predicted for those regions. (B) Coomassie Blue-stained SDS-PAGE gels of sfGFP-DCLK1 proteins used in this study.

**Figure S2.**
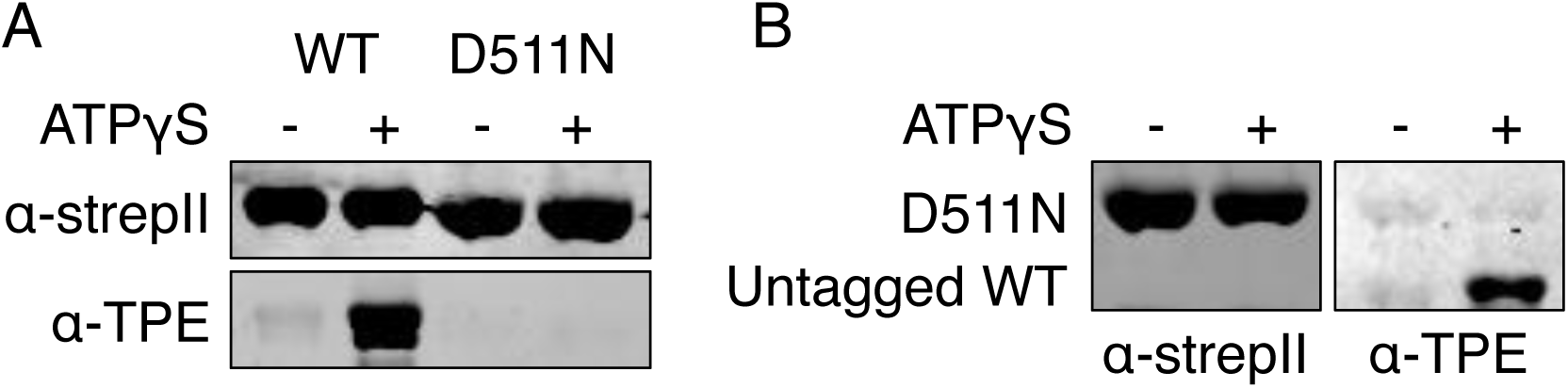
DCLK1 autophosphorylates via an intramolecular mechanism. Related to Figure 1. (A) Immunoblots of strepII-sfGFP-tagged WT or kinase dead (D511N) DCLK1 incubated in the absence or presence of ATPγS for 30 min at 37°C. DCLK1-WT, but not DCLK1-D511N, robustly autophosphorylates. (B) Immunoblots of strepII-sfGFP-tagged DCLK1-D511N incubated with an untagged version of DCLK1-WT to distinguish the proteins by size in the absence or presence of ATPγS for 30 min at 37°C. While DCLK1-WT autophosphorylates, there is still no detectable level of phosphorylation for DCLK1-D511N. For *A* and *B*, primary antibodies were mouse anti-strep (Fisher NBP243719) and rabbit anti-thiophosphate ester (Abcam ab133473).

**Figure S3.**
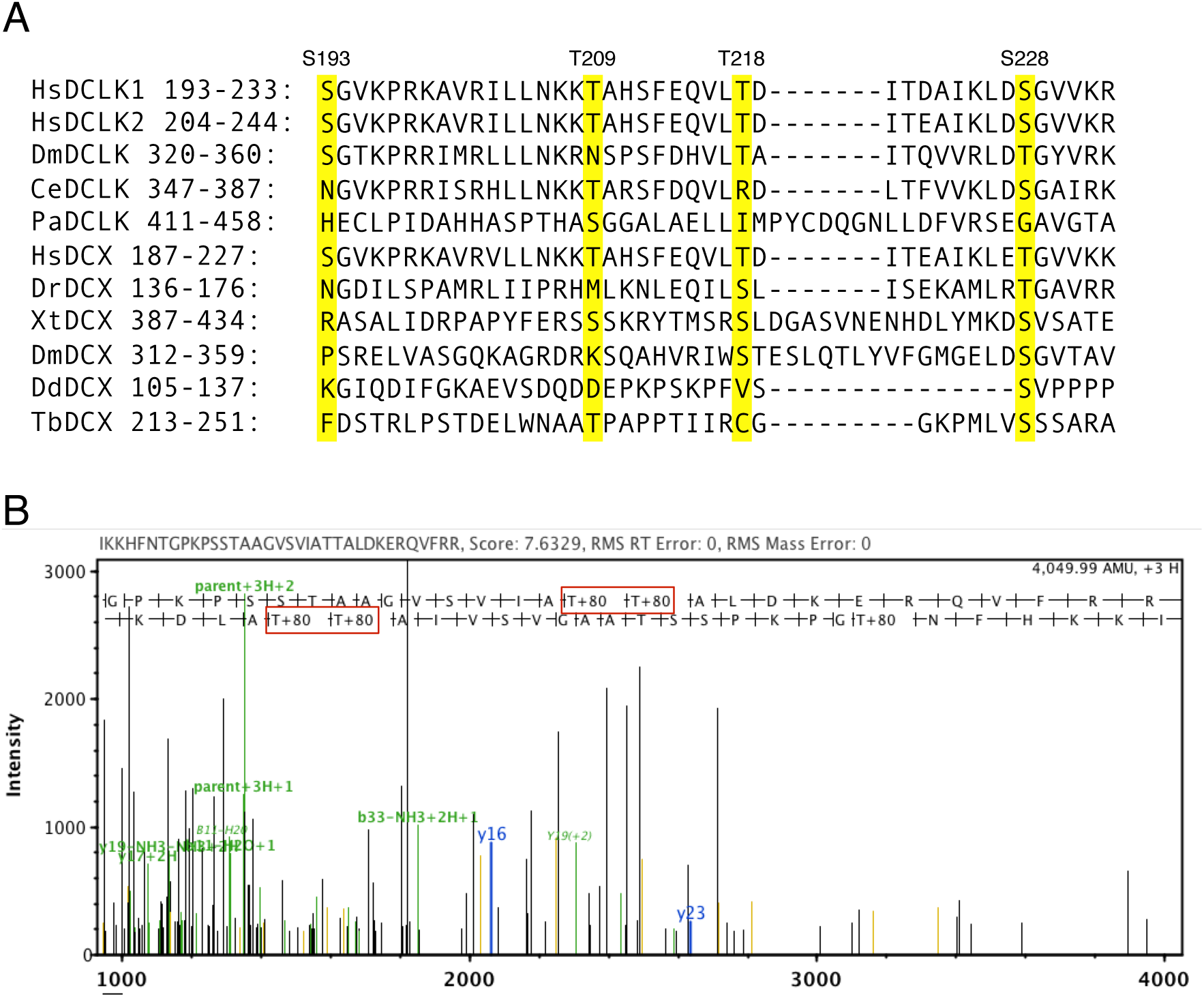
Dissection of the phosphorylated residues within DCLK1. Related to Figures 3-5. (A) Sequence alignment of the region of DC2 in DCLK or DCX including the four amino acid residues mutated in ΔC_4A_ (highlighted in yellow). Hs: Homo sapiens, Dm: Drosophila melanogaster, Ce: Caenorhabditis elegans, Pa: Pseudozyma aphidis, Dr: Danio rerio, Xt: Xenopus tropicalis, Dd: Dictyostelium discoideum, and Tb: Trypanosoma brucei. (B) Representative ion spectrum of a DCLK1-WT peptide containing phosphorylation at T687 and T688 detected by LC-MS/MS. The major y and b ions are labeled in the spectrum and the phosphorylated amino acids are boxed in red.

**Figure S4.**
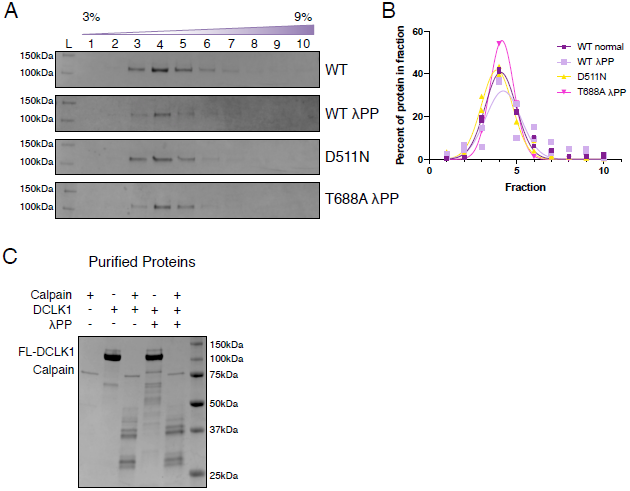
DCLK1 autophosphorylation does not grossly alter protein conformation or cleavage. Related to Figures 3-5. (A) Coomassie Blue-stained SDS-PAGE gels of purified DCLK1 proteins fractionated by sucrose density gradient centrifugation. Representative gels from n = 2 independent experiments for each protein except n = 1 for T688A. (B) Quantification of the average percent of DCLK1 protein in each fraction, revealing a peak in fraction 4 for all DCLK1 proteins. (C) Coomassie Blue-stained SDS-PAGE gel of purified DCLK1-WT protein expressed in bacteria under standard conditions (lanes 2 and 3) and DCLK1-WT protein co-expressed with λPP in bacteria (lanes 4 and 5) incubated with calpain or incubated in buffer alone for 15 min at 30°C show similar band patterns after cleavage with calpain.

